# Brain age prediction reveals aberrant brain white matter in schizophrenia and bipolar disorder: A multi-sample diffusion tensor imaging study

**DOI:** 10.1101/607754

**Authors:** Siren Tønnesen, Tobias Kaufmann, Ann-Marie de Lange, Genevieve Richard, Nhat Trung Doan, Dag Alnæs, Dennis van der Meer, Jaroslav Rokicki, Torgeir Moberget, Ivan I. Maximov, Ingrid Agartz, Sofie R. Aminoff, Dani Beck, Deanna Barch, Justyna Beresniewicz, Simon Cervenka, Helena Fatouros Bergman, Alexander R. Craven, Lena Flyckt, Tiril P. Gurholt, Unn K. Haukvik, Kenneth Hugdahl, Erik Johnsen, Erik G. Jönsson, KaSP^i^, Knut K. Kolskår, Kristiina Kompus, Rune Andreas Kroken, Trine V. Lagerberg, Else-Marie Løberg, Jan Egil Nordvik, Anne-Marthe Sanders, Kristine Ulrichsen, Ole A. Andreassen, Lars T. Westlye

## Abstract

**Background:** Schizophrenia (SZ) and bipolar disorders (BD) share substantial neurodevelopmental components affecting brain maturation and architecture. This necessitates a dynamic lifespan perspective in which brain aberrations are inferred from deviations from expected lifespan trajectories. We applied machine learning to diffusion tensor imaging (DTI) indices of white matter structure and organization to estimate and compare brain age between patients with SZ, BD, and healthy controls across 10 cohorts.

**Methods:** We trained six cross-validated models using different combinations of DTI data from 927 healthy controls (HC, 18-94 years), and applied the models to the test sets including 648 SZ (18-66 years) patients, 185 BD patients (18-64 years), and 990 HC (17-68 years), estimating brain age for each participant. Group differences were assessed using linear models, accounting for age, sex, and scanner. A meta-analytic framework was applied to assess the heterogeneity and generalizability of the results.

**Results:** 10-fold cross-validation revealed high accuracy for all models. Compared to controls, the model including all feature sets significantly over-estimated the age of patients with SZ (*d*=-.29) and BD (*d*=.18), with similar effects for the other models. The meta-analysis converged on the same findings. Fractional anisotropy (FA) based models showed larger group differences than the models based on other DTI-derived metrics.

**Conclusions:** Brain age prediction based on DTI provides informative and robust proxies for brain white matter integrity. Our results further suggest that white matter aberrations in SZ and BD primarily consist of anatomically distributed deviations from expected lifespan trajectories that generalize across cohorts and scanners.

## Introduction

Schizophrenia (SZ) and bipolar (BD) spectrum disorders are severe mental disorders with partly overlapping clinical characteristics and pathophysiology. Both are highly heritable (1) with a substantial neurodevelopmental aetiology (2, 3). Along with evidence of accelerated age-related brain changes in adult patients with SZ (4-6), the neurodevelopmental origin supports a dynamic lifespan perspective in which genetic and biological factors interact with age-related environmental and physiological processes.

Aberrant myelination and brain wiring during adolescence has been included among the neurobiological features of severe mental disorders, and white matter (WM) aberrations have been documented prior to disease onset (7-11). Brain imaging has shown that normative WM development follows a characteristic non-linear trajectory with peak maturation around the third or fourth decade (12-14). Compared to healthy controls (HC), adult patients with SZ or BD exhibit anatomically distributed group-level differences in various diffusion-based indices of WM structure (15, 16).

Supporting a neurodevelopmental origin, it has been demonstrated that patients with adolescent-onset SZ show WM aberrations (17), and that their developmental trajectory is altered and delayed (18) compared to age-matched, normally developing peers. Further, children and adolescents with increased symptom burden, albeit presumably at subclinical levels, were found to exhibit altered diffusion-based WM properties compared to peers with low or no symptoms of mental distress (19), highlighting a critical role of WM development in mental health in youths. To which degree group differences observed between adult patients and HC accelerate during the course of the adult lifespan is unclear. The neurodegenerative account of schizophrenia and severe mental illness is debated (20) and lacks unequivocal support from imaging studies (16, 21), but some studies have suggested stronger age-related deterioration of the brain in patients compared to HC (22, 23).

Despite converging evidence of case-control differences both preceding and following disease onset, recent brain imaging studies have documented substantial heterogeneity within patient groups (24, 25). In contrast to conventional group-level analyses, brain age prediction using machine learning on imaging features allows for brain-based phenotyping at the individual level, and enables an efficient dimensionality reduction of the neuroimaging data into one or more biologically informative summary measures (26, 27). The discrepancy between an individual’s chronological age and predicted brain age, referred to as the *brain age gap* (BAG), has been found to be higher in patients with SZ (5, 28, 29) and several other brain disorders (29). However, these previous studies have exclusively used brain grey matter features for brain age prediction. Thus, given the well-documented role of WM aberrations in patients with mental illness (15, 30-32), brain age prediction based on diffusion imaging is clearly warranted.

In order to fill this current gap in the literature, we here compared individual BAGs between patients diagnosed with SZ or BD and HC, using four conventional metrics (fractional anisotropy (FA), mean diffusivity (MD), radial diffusivity (RD) and axial diffusivity (AD)) obtained from diffusion tensor imaging (DTI). We used an independent training set comprising 927 HC aged 18-94 years, and applied the resulting model to our test sample including patients with SZ (n=648), and BD (n=185), as well as HC (n=990), from 10 independent cohorts (see Materials and Methods for details). In order to specifically assess the robustness and quantify the heterogeneity of effects across cohorts, we adopted a meta-analytic statistical framework in addition to a mega-analysis across cohorts.

We trained six different models based on various combinations of the DTI metrics, which allowed us to compare prediction accuracy and subsequent group differences for each model. Based on converging evidence of widespread WM aberrations in patients with severe mental disorders (15), we hypothesized higher BAG in patients with SZ and BD compared to HC, with stronger effects in SZ compared to BD. To test the relevance of the varying spatial resolution of the feature sets, which is important to inform the discussion regarding the anatomical specificity of brain WM aberrations, we compared models including various atlas-based tracts of interest (TOIs) with models including only global features. Based on previous studies comparing the age prediction accuracy and clinical sensitivity between metrics (16, 27, 33), we hypothesized high age prediction accuracy and sensitivity to group differences, but remained agnostic concerning the relative ranking of the various features.

## Materials and methods

We combined diffusion MRI data from 2750 individuals from 11 sites/studies across 10 different scanners. Figure 1A, Supplemental Figures 1-2, and Supplemental Tables 1-2 summarise key demographics for each cohort. Supplemental Table 3 summarizes the MRI systems and diffusion acquisition protocols.

**Figure 1.**
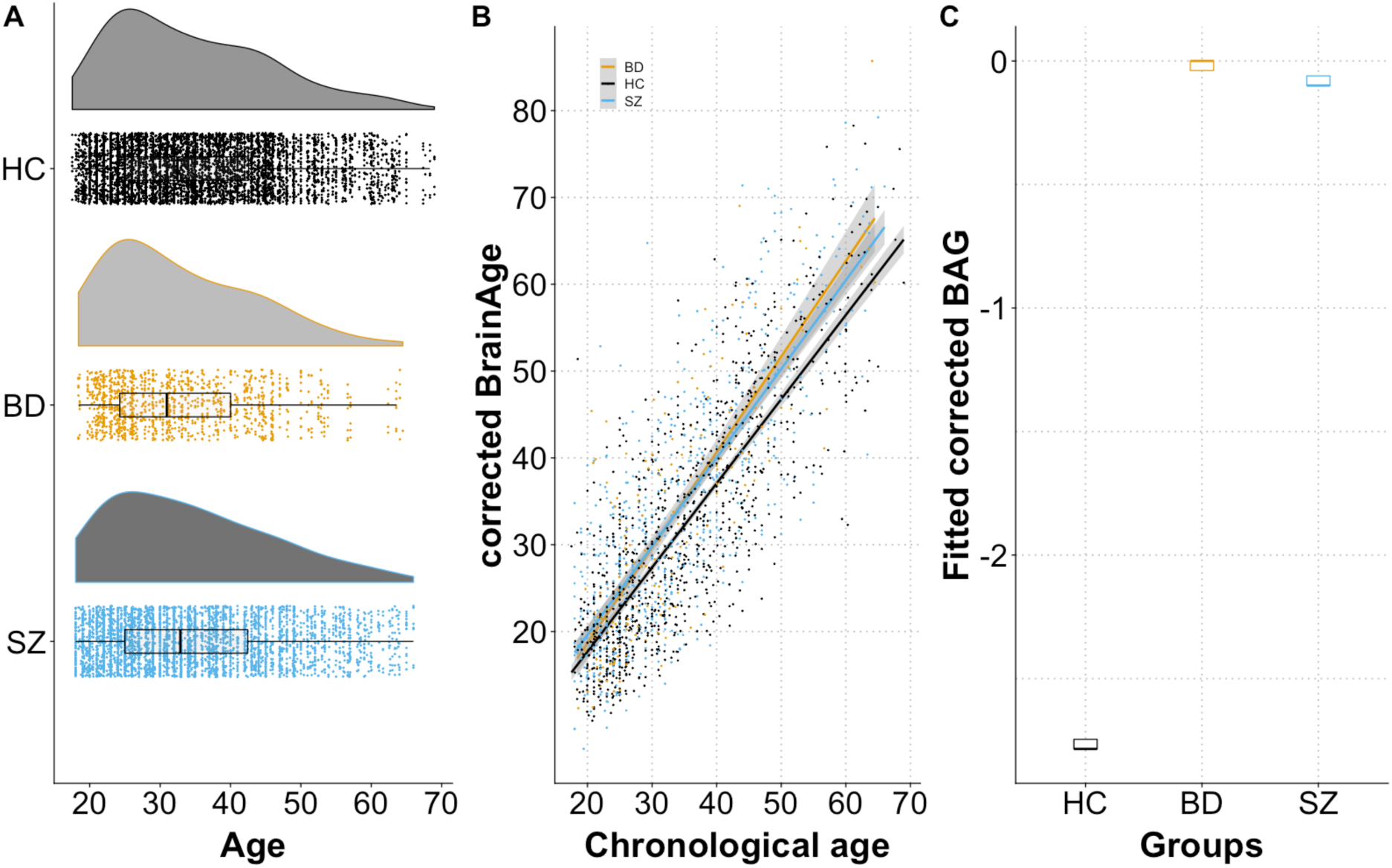
A) Raincloud plot depicting the age distribution for each diagnostic group in the test sets. Density plots are shown on top with data points and boxplot underneath. B) Predicted age (corrected for age and scanner) based on the all features model plotted as function of chronological age. The fit lines represent the best linear fit within each group. C) Boxplots plots showing the distribution of fitted corrected BAG in each group.

The dataset was split into a training set and a test set. Supplemental Figure 2 shows the age distribution within each of the two cohorts in the training set. Briefly, the training set consisted of 927 HC covering the full adult lifespan (mean age=53.81 years, s.d.=18.38, range 18-94 years). The test set comprised 990 HC (mean age=34.70 years, s.d.=11.24, range=17-68), 185 patients with BD (mean age=33.12 years, s.d.=10.53, range=18-64), and 648 patients with SZ (mean age=34.49 years, s.d.=11.40, range=18-66).

**Figure 2.**
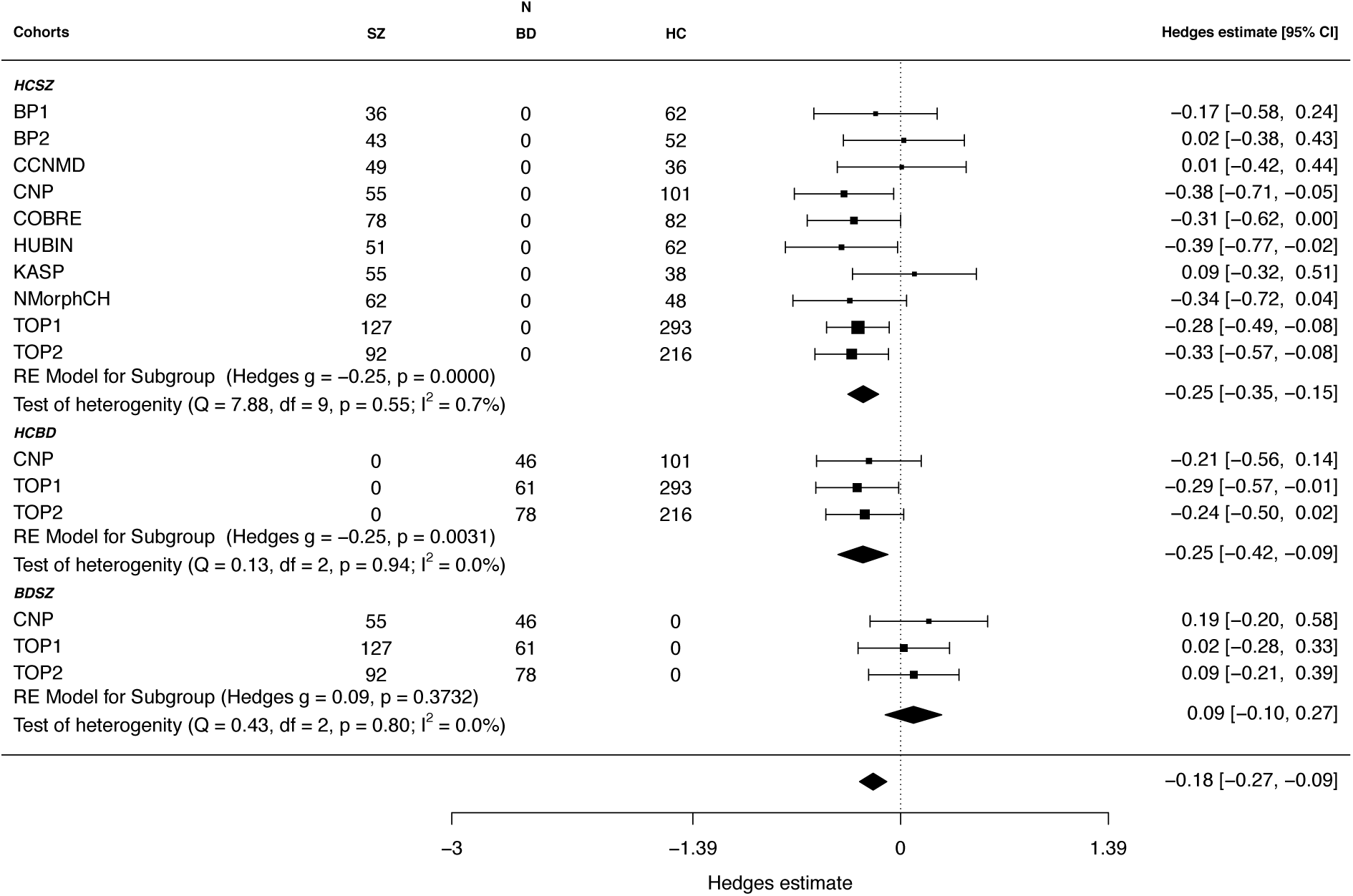
Forest plot summarizing the results from the meta-analytical approach for the all features model. Hedges (*g*) estimate was used to calculate the effect size.

### MRI acquisition and processing

A summary of MRI acquisition protocol for each cohort is presented in Supplementary Table S3. Imaging analyses were performed using the Oxford Center for Functional Magnetic Resonance Imaging of the Brain (FMRIB) Software Library (FSL) (34-36). To correct for signal loss, motion and eddy currents all cohorts were processed using eddy (http://fsl.fmrib.ox.ac.uk/fsl/fslwiki/eddy) (37, 38). Two cohorts (TOP1 and TOP2) had collected blip-up/blip-down sequences, and were additionally processed using topup (http://fsl.fmrib.ox.ac.uk/fsl/fslwiki/topup) (34, 39) prior to eddy. Using an integrated framework along with correction for susceptibility-induced distortions, eddy currents and motion, eddy detects and replaces slices affected by signal loss due to bulk motion during diffusion encoding (38).

Fitting of the diffusion tensor was done using dtifit in FSL, yielding conventional DTI metrics, including fractional anisotropy (FA), and mean (MD), radial (RD) and axial (AD) diffusivity. FA, MD, RD and AD maps were further processed using tract-based spatial statistics (TBSS) (40). FA volumes were skull-stripped and aligned to the FMRIB58_FA template supplied by FSL using nonlinear registration (FNIRT) (41). Next, mean FA were derived and thinned to create a mean FA skeleton, representing the center of all tracts common across subjects. We thresholded and binarized the mean FA skeleton at FA>0.2. The procedure was repeated for MD, AD, and RD. For each individual, we calculated the mean skeleton value for each metric, as well as mean values within 23 TOIs (Supplemental Table 4) based on two probabilistic white matter atlases (CBM-DTI-81 white-matter labels atlas and the JHU white-matter tractography atlas (42-44)). In total, we derived 96 DTI features per individual including the mean skeleton values.

### Quality assessment

Subjects with poor image quality due to subject motion or other visible image artefacts (e.g. due to metal) were removed (n=160, including HC (n=59), patients with SZ (n=39), patients with BD (n=28), and individuals with missing information (n=34)). Demographics of the excluded participants are presented in Supplemental Table 5. Additionally, we employed a multistep quality assessment (QA) procedure (16) that included maximum voxel intensity outlier count (MAXVOX) and tSNR (45) prior to statistical analyses. Briefly, we ran the QA iteratively excluding participants with a QA score of 2.5 SD below the mean. In order to compute the QA score we inverted the MAXVOX score, z-normalized both scores independently (MAXVOX and tSNR), and computed a summary score combing the two scores. In short, manual inspection of the flagged datasets after QA suggested adequate quality. Thus, we present results on the full dataset with supplemental results from a stringent QA (see (16) for additional information).

### Brain age prediction

We trained six age prediction models. Our main model included all 96 features across all DTI metrics. To assess sensitivity for each metric separately, we trained four additional models based on all TOIs for each metric (FA, RD, MD or AD). To test the value of including regionally specific information, we trained an additional model with only the global mean skeleton feature from all four metrics included.

The following pipeline for brain age prediction was identical for all six models: We used the xgboost framework in R (46) to build the prediction model. The number of rounds (nround), maximum depth (max_depth) and subsample were tuned and optimised using a 5-fold cross validation of the training data, with early stopping if the prediction errors did not improve for 20 rounds. The learning rate (eta) was pre-set to eta=0.01. Besides the default setting, the following parameters were used in the model: nround=1400, max_depth=14.

Prior to implementing the model, we regressed out the main effect of scanner from the DTI features in the entire dataset while accounting for age, age^2^, and sex using linear models in R (47). To estimate the reliability of our age prediction model, we used a 10-fold cross-validation procedure within the training sample and repeated the cross-validation step 100 times to provide a robust estimate of model prediction. Within the same procedure, we tested the performance of our trained model by predicting age in unseen subjects in the test sample. By applying the model to the test sample 100 times we obtain both a mean estimate and an estimate of uncertainty. For each iteration we calculated BAG defined as the difference between chronological and predicted age. For each individual in the test set we computed the average BAG across the 100 folds and corrected these values for main effects of scanner and a well-documented age-related bias using linear models, per previous recommendations (48). Next, based on the age- and scanner-corrected BAG we computed a corrected brain age for each individual, and then computed the mean absolute error (MAE), the root mean squared error (RMSE) and the correlation between corrected predicted age and chronological age as measures of model performance.

### Statistical analyses

Statistical analyses were performed using R (version 3.3.3 (2017-03-06)(47)). We tested for main effects of diagnosis using linear models with corrected BAG as dependent variable and group, sex, age and scanner site as independent variables, and performed pairwise group comparisons as appropriate. Using the metafor package (49) in R we adopted a meta-analytic framework in order to assess the heterogeneity and generalizability of the results. A random-effects model was used to weigh the primary studies prior to aggregating the effect size. Effect sizes were aggregated using the estimated marginal means of BAG from each group contrast (HC/SZ, HC/BD and BD/SZ), accounting for age, age^2 and sex. For effect size estimates, we used Hedges’ *g*. Cochran’s heterogeneity statistic *Q* was used to test the homogeneity of effect sizes. A χ^2^ test with *k*-1 degrees of freedom was used to examine the significance of Cochran’s *Q.* The heterogeneity was quantified using the *I*^2^ statistic, which is sensitive to the degree of inconsistency in results between cohorts.

## Results

### Brain age predictions

Age prediction in the *training* set using 10-fold cross validation revealed high correlations between chronological and predicted age for the main model including all features (*r*=.924, 95% CI: .912-.935, MAE=6.49, RMSE=8.08, Supplemental Figure 3).

Figure 1B shows predicted age plotted as a function of chronological age for the unseen test set when using the full feature set, and Table 1 summarizes the prediction accuracy for all six models. The age prediction models generalized to HC (r=.806, MAE=6.92, RMSE=8.46), patients with BD (r=.808, MAE=6.85, RMSE=8.50), and patients with SZ (r=.798, MAE= 7.11, RMSE=8.79). While all models performed relatively well, prediction accuracy was highest for the full model, and the global mean skeleton model outperformed the ROI based single-metrics models. Supplemental Figure 4 shows the correlation matrix between all models, indicating a strong correlation between all models with the exception of FA and AD.

**Table 1.** Results from the six brain age models.

| | Correlation<br>between<br>chronological and<br>predicted age<br>(MAE/RMSE <sup>a</sup>) | Corrected<br>BAG SZ<br>Mean ±sd<br>(95%CI) | Corrected<br>BAG BD<br>Mean ±sd<br>(95%CI) | Corrected<br>BAG HC<br>Mean ±sd<br>(95%CI) | F | $p^b$ | Pairwise<br>comparis<br>on | Cohens d<br>HC/SZ,<br>BD/HC, BD/SZ | Cohens d<sup>c</sup><br>Mean skeleton<br>(SZ/HC, BD/HC,<br>BD/SZ) |
| --- | --- | --- | --- | --- | --- | --- | --- | --- | --- |
| All features | .80 (6.98/8.58) | -0.09±8.79<br>[-0.77,0.59] | -0.02±8.53<br>[-1.26,1.22] | -2.77±8.00<br>[-3.27,-2.27] | 23.01 | <.001 | HC<SZ,<br>HC<BD | -.29/.18/-.03 | |
| Mean<br>skeleton | .76 (7.91/9.75) | 0.56±10.04<br>[-0.21,1.34] | -0.16±8.97<br>[-1.46,1.14] | -2.55±9.37<br>[-3.13,-1.97] | 21.57 | <.001 | HC<SZ,<br>HC<BD | .30/.14/-.04 | |
| FA | .77 (7.61/9.50) | 0.24 ±9.92 [-<br>0.52,1.01] | 0.21±9.42<br>[-1.16,1.57] | -3.23±8.66<br>[-3.77,-2.69] | 30.55 | <.001 | HC<SZ,<br>HC<BD | -.33 /.21/ <.01 | -.36/-.17/.18 |
| RD | .78 (7.65/9.32) | -0.60±9.72<br>[-1.35,0.15] | -0.05±8.64<br>[-1.30,1.20] | -3.45±8.50<br>[-3.98,-2.92] | 24.59 | <.001 | HC<SZ,<br>HC<BD | -.29/.21/-.03 | .35/.20/-.14 |
| MD | .78 (7.54/9.23) | -1.33±9.40<br>[-2.07,-0.61] | -0.21±8.49<br>[-1.44,1.02] | -3.45±8.54<br>[-3.98,-2.92] | 17.27 | <.001 | HC<SZ,<br>HC<BD | .22/.20/.06 | .24/.17/-.06 |
| AD | .80 (7.15/8.80) | -2.17±8.46<br>[-2.82,-1.52] | -0.46±8.38<br>[0.75,-1.67] | -2.95±8.43<br>[-3.48,-2.42] | 6.95 | .001 | HC>BD | -.10/.16/.10 | <.001/.08/.09 |

Abbreviations: MAE: Mean absolute error, RMSE: Root mean squared error, FA: fractional anisotropy, MD: mean diffusivity, RD: radial diffusivity and AD: axial diffusivity.
<sup>a</sup>Correlations, MAE and RMSE are reported for predicted age corrected for scanner and age.
<sup>b</sup> $\alpha$ level for Bonferroni-corrected $p$ -values = .008)
<sup>c</sup>Cohens d reported for the pairwise group comparisons for mean skeleton DTI values

### Group differences in BAGR

Table 1 and Supplemental Figure 5 summarize the results from the group comparisons from the six models and Figure 1C shows the distributions of fitted corrected BAG within each group for the all feature model. Briefly, all models revealed significant main effects of group, with higher corrected BAG in patients with SZ and BD compared to HC, with effect sizes ranging between d=-0.10 and d=0.33. The model based on FA yielded the strongest effect size for the main group effect, although the models including MD and RD in addition to FA revealed similar patterns. The model based on AD revealed less consistent results, and was the only model not showing significant group differences between SZ and HC.

### Meta-analysis and heterogeneity in effects between cohorts

Figure 2 shows a forest plot summarizing the results from the meta-analysis for BAG computed using the full feature model. Supplemental Figures 6-10 show the results from the other models. In short, in line with the mega-analysis the results revealed significantly higher BAG in SZ and BD compared to HC, with moderate effect sizes. The analysis did not support a group difference in brain age gap between BD and SZ. Whereas the effect sizes varied slightly between cohorts for the full model, the *Q* and *I*^2^ statistics indicated low and non-significant heterogeneity. Supplemental Figure 11 shows each cohort’s contribution to the heterogeneity and influence on the result from the meta-analysis.

### Quality control

Supplemental Figure 12 summarises the results from multistep QA. Briefly, higher corrected BAG was observed in SZ and BD compared to HC across all levels of QA, with highly similar effect sizes.

## Discussion

The aetiology of severe mental disorders has a substantial neurodevelopmental component, which is amongst other characteristics reflected in altered brain maturational trajectories during the formative years of childhood and adolescence, and as group-level differences in adult patient populations. Along with evidence of genetic and clinical overlap with several aging-related conditions, including cardiovascular risk factors and increased mortality, the neurodevelopmental account supports the need for a dynamic lifespan perspective in the search for disease mechanisms. Here, in ten different cohorts comprising healthy controls and patients with SZ and BD, we used machine learning to estimate brain age using DTI indices of white matter structure and organization. This novel approach yielded five main results. First, in a large independent training set, we found high accuracy of brain age prediction across the adult lifespan using DTI features, which largely generalized to the independent test set, supporting the feasibility and sensitivity of the approach. Second, applying the model to an independent test set revealed significantly higher brain age gap in patients with SZ and BD compared to HC. Third, follow-up meta-analysis and tests of heterogeneity suggested high consistency across independent cohorts and scanners. Fourth, brain age models based on FA showed higher sensitivity than models based on the other metrics, both alone and combined. Finally, the reduced set of global mean skeleton features compared to a number of regional atlas-based features revealed highly converging results. We next discuss the implications of these findings in more detail.

Brain age prediction provides an informative summary measure that may serve as a proxy for brain integrity and health across normative and clinical populations. Neuroimaging derived white and grey matter phenotypes carry distinct biological information of brain integrity, and tissue-specific brain age models may provide higher sensitivity and specificity to relevant biological processes compared to conventional models based on grey matter features alone (27). DTI has been broadly applied in clinical neuroscience due to its proposed sensitivity to microstructural properties of brain tissue. However, whereas previous studies have documented higher brain age in patients with severe mental disorders, these were based on grey matter models only (5, 28, 29). In order to test if previous findings suggesting clinical deviations from normative grey matter trajectories generalize to white matter, we performed brain age prediction using different combinations of DTI metrics. In line with previous brain age prediction studies using diffusion MRI (27, 50) we obtained high age prediction accuracy across most models. In accordance with previous evidence suggesting that regional DTI-based indices of brain aging reflect relatively low-dimensional and global processes (12, 51), we found similar prediction accuracy for the reduced models comprising global mean skeleton values and the models including extended sets of regional features. Although brain white matter aging shows some regional heterogeneity (12), these findings demonstrate that the most relevant information required for brain age prediction is captured at a global level. This conjecture is also supported by a recent twin study demonstrating that a large proportion of the estimated heritability of specific tracts is accounted for by a general factor (52).

Likewise, we found that the sensitivity to group differences was not strongly dependent on the inclusion of the full feature set. Indeed, the effect size obtained when comparing patients with SZ and HC were slightly higher for the global mean skeleton model compared to the full model. These findings are in line with recent evidence of anatomically widely distributed group differences between healthy controls and patients with SZ (15). Interestingly, the largest effect when comparing SZ and HC was obtained for the FA only model, supporting the sensitivity of FA to clinical differences in WM properties (15, 16). Higher predicted brain age in the patient groups compared to healthy controls may indicate altered rate of brain maturation or accelerated brain aging in patients with severe mental disorders. However, our cross-sectional design does not permit us to make inference about brain development or aging per se, and previous reports of relatively age-invariant group differences in brain volumetry (21) and DTI indices (16) suggest that the reported group differences in brain age may reflect differences accumulating early in life. Unfortunately, due to the current study design with adults only we cannot address the maturational trajectories in the formative years. Although the application of diffusion MRI as the basis for age prediction is novel, higher gray matter brain age has been shown in several brain and mental disorders (29, 53). We expand these previous findings by documenting higher DTI based white matter brain age in both SZ and BD, and, although with moderate effect sizes, we show that the effects generalize relatively well across cohorts and scanners, with only minor heterogeneity in effect sizes between cohorts.

We found no significant difference in DTI based brain age between BD and SZ, supporting previous evidence of partly overlapping clinical and biological characteristics between these two diagnostic categories (16, 54, 55). While the current results support the existence of a common set of mechanisms across disorders, future studies utilizing a broader range of imaging modalities in combination with specific genetic, clinical, cognitive, sociodemographic and biological phenotypes may allow for the identification of specific diagnostic signatures and sub-groups. However, inherent limitations associated with the classical case-control design in mental health research have recently been emphasized using neuroimaging data (24, 25). In particular, the current lack of biologically informed diagnostic criteria should motivate future studies to consider alternative approaches to promote a novel clinical nosology based both on symptomatology and data-driven clustering (56), as well as brain-based and biological phenotypes cutting across diagnostic boundaries.

Our results document robust group-level deviances in white matter structure manifesting as older-appearing brains in patients with severe mental disorders compared to their healthy peers. Whereas DTI-based markers are sensitive to different biological and anatomical characteristics, the current specificity does not allow for inference on the distinct neurobiological mechanisms involved. Myelin integrity and myelin packing density are among the proposed candidate mechanisms for observed changes in DTI metrics (57-59), but the specificity is low, and the current results probably reflect a combination of neurobiological processes and macroanatomical differences. Previous evidence implicated myelin-related abnormalities and neuroinflammation both in the pathophysiology of severe mental disorders and in brain aging (60-63). Future studies may benefit from the inclusion of advanced multi-shell diffusion MRI allowing for stronger inference on the microstructural milieu of the brain tissue, including microstructural indices based on different diffusion scalar metrics (e.g., Neurite Orientation Dispersion and Density Imaging (NODDI) (64, 65), diffusion kurtosis imaging (DKI) (66), white matter tract integrity (WMTI) (67) and restriction spectrum imaging (RSI) (68)).

In line with previous findings of widely distributed effects in well-powered studies of brain aging (12) and schizophrenia (15), we found similar age prediction accuracy and subsequent group differences in brain age for the model including only global mean skeleton values and the model including a range of regionally informative values extracted from various atlas-based tracts and regions of interest. Although specific symptoms and clinical traits may map preferentially onto specific neuroanatomical subsystems (see e.g. (19)), these novel results suggest that a large proportion of the variance associated with age and corresponding deviations in the patient groups are captured by primarily global brain processes, with relevance for our understanding of the anatomical heterogeneity and dimensionality of brain aging and severe mental illness.

In addition to the anatomical distribution of effects, the spatiotemporal dynamics of brain development and aging and their deviations in patients with mental disorders remain unclear. The individual level onset and rate of the group-level deviations from the normative white matter trajectory is unknown and can only be inferred using longitudinal designs covering sensitive periods of neurodevelopment. Previous studies have shown both delayed neurodevelopment during adolescence (18) and accelerated aging in adulthood (5) in patients with severe mental disorders. Whereas these observations are not mutually exclusive, future studies should aim at disentangling the lifespan dynamics, e.g. by including individuals with a wider age range, and pursuing longitudinal designs including individuals across a wide range of functional levels and risk. The latter may be particularly pertinent to disentangle primary disease-related mechanisms and secondary factors related to the disease, including medication and life-style factors such as nutrition, physical activity, education, and a range of sociodemographic variables, which all interact with key neurodevelopmental processes (69) Unfortunately, although possible effects of psychotropic drugs on the brain is a topic of great interest and importance (70-72), in common with other studies employing a cross-sectional and non-randomised design, the current design does not allow us to make inference about the effects of medication and other clinical and lifestyle factors on brain age, which should be investigated by future and properly designed studies. Meanwhile, previous studies reporting associations with medication status in smaller samples need to be interpreted in light of the recent lack of significant associations in the largest DTI study to date (15). We did not exclude white matter hyperintensities (WMH) in the training or test sets, and future studies including a wider range of MRI modalities are necessary to determine the possible confounding effects of WMH on the age prediction models and subsequent group comparisons.

In conclusion, in this multi-sample study including patients from 10 different cohorts, we report higher brain age in patients with SZ and BD compared to HC using various DTI-based indices of white matter structure and organization. In contrast to most previous studies comparing diffusion MRI metrics directly between groups we used a multi-sample approach, which allowed us to specifically assess generalizability across 9-10 different cohorts, sites and scanners. These results represent a highly relevant contribution the field and an important supplement to previous reports, which have largely ignored between-sample heterogeneity and generalizability. Although the effect sizes were modest, our unique design allowed us to specifically quantify the heterogeneity and robustness of effects across cohorts and scanners, supporting that brain age prediction using diffusion MRI is a sensitive marker in the clinical neurosciences.

## Acknowledgements

This work was funded by the South-Eastern Norway Regional Health Authority (2014097, 2015073, 2016083, 2016044, 2019101), the Western Norway Regional Health Authority (911820, 911679), the Research Council of Norway (213700, 204966, 249795, 223273, 213727, 298646, 300768), KG Jebsen Stiftelsen, the European Commission’s 7th Framework Programme (#602450, IMAGEMEND), and the European Research Council under the European Union’s Horizon 2020 research and Innovation program (ERC StG, Grant 802998). KaSP was supported by grants from the Swedish Medical Research Council (SE: 2009-7053; 2013-2838; SC: 523-2014-3467), the Swedish Brain Foundation, Åhlén-siftelsen, Svenska Läkaresällskapet, Petrus och Augusta Hedlunds Stiftelse, Torsten Söderbergs Stiftelse, the AstraZeneca-Karolinska Institutet Joint Research Program in Translational Science, Söderbergs Königska Stiftelse, Professor Bror Gadelius Minne, Knut och Alice Wallenbergs stiftelse, Stockholm County Council (ALF and PPG), Centre for Psychiatry Research, KID-funding from the Karolinska Institutet. Data collection and sharing for this project was provided by the Cambridge Centre for Ageing and Neuroscience (CamCAN). CamCAN funding was provided by the UK Biotechnology and Biological Sciences Research Council (grant number BB/H008217/1), together with support from the UK Medical Research Council and University of Cambridge, UK. CCNMD was supported through NIH Grants P50 MH071616 and R01 MH56584. CNP was supported by the Consortium for Neuropsychiatric Phenomics (NIH Roadmap for Medical Research grants UL1-DE019580, RL1MH083268, RL1MH083269, RL1DA024853, RL1MH083270, RL1LM009833, PL1MH083271, and PL1NS062410). The HUBIN project was supported by the Swedish Research Council (2006-2992, 2006-986, K2007-62X-15077-04-1, 2008-2167, K2008-62P-20597-01-3. K2010-62X-15078-07-2, K2012-61X-15078-09-3, 2017-00949, K2015-62X-15077-12-3), the regional agreement on medical training and clinical research between Stockholm County Council and the Karolinska Institutet, the Knut and Alice Wallenberg Foundation. StrokeMRI was supported by the Research Council of Norway (249795, 248238), the South-Eastern Norway Regional Health Authority (2014097, 2015044, 2015073, 2016083), and the Norwegian ExtraFoundation for Health and Rehabilitation (2015/FO5146). Data collection and sharing for the NMorphCH project was funded by NIMH grant A R01 MH056584. The BergenPsykose project was supported by the European Research Council (ERC AdG) (693124), and Western Norway Health-Authorities (912045). A pre-print version of the manuscript was published in BioRxiv.

^i^ Members of the Karolinska Schizophrenia Project (KaSP): Farde L^1^, Flyckt L^1^, Engberg G^2^, Erhardt S^2^, Fatouros-Bergman H^1^, Cervenka S^1^, Schwieler L^2^, Piehl F^3^, Agartz I^1,4,5^, Collste K^1^, Victorsson P^1^, Malmqvist A^2^, Hedberg M^2^, Orhan F^2^, Sellgren C^2^

^1^Centre for Psychiatry Research, Department of Clinical Neuroscience, Karolinska Institutet, & Stockholm County Council, Stockholm, Sweden; ^2^Department of Physiology and Pharmacology, Karolinska Institutet, Stockholm, Sweden; ^3^Neuroimmunology Unit, Department of Clinical Neuroscience, Karolinska Institutet, Stockholm, Sweden; ^4^NORMENT, Division of Mental Health and Addiction, University of Oslo, Oslo, Norway; ^5^Department of Psychiatry Research, Diakonhjemmet Hospital, Oslo, Norway

