## Supplemental material for "Brain age prediction reveals aberrant brain white matter in schizophrenia and bipolar disorder: A multi-sample diffusion tensor imaging study"

Short title: White matter brain age in severe mental illness

Keywords: brain age, DTI, psychosis, schizophrenia, bipolar, machine learning

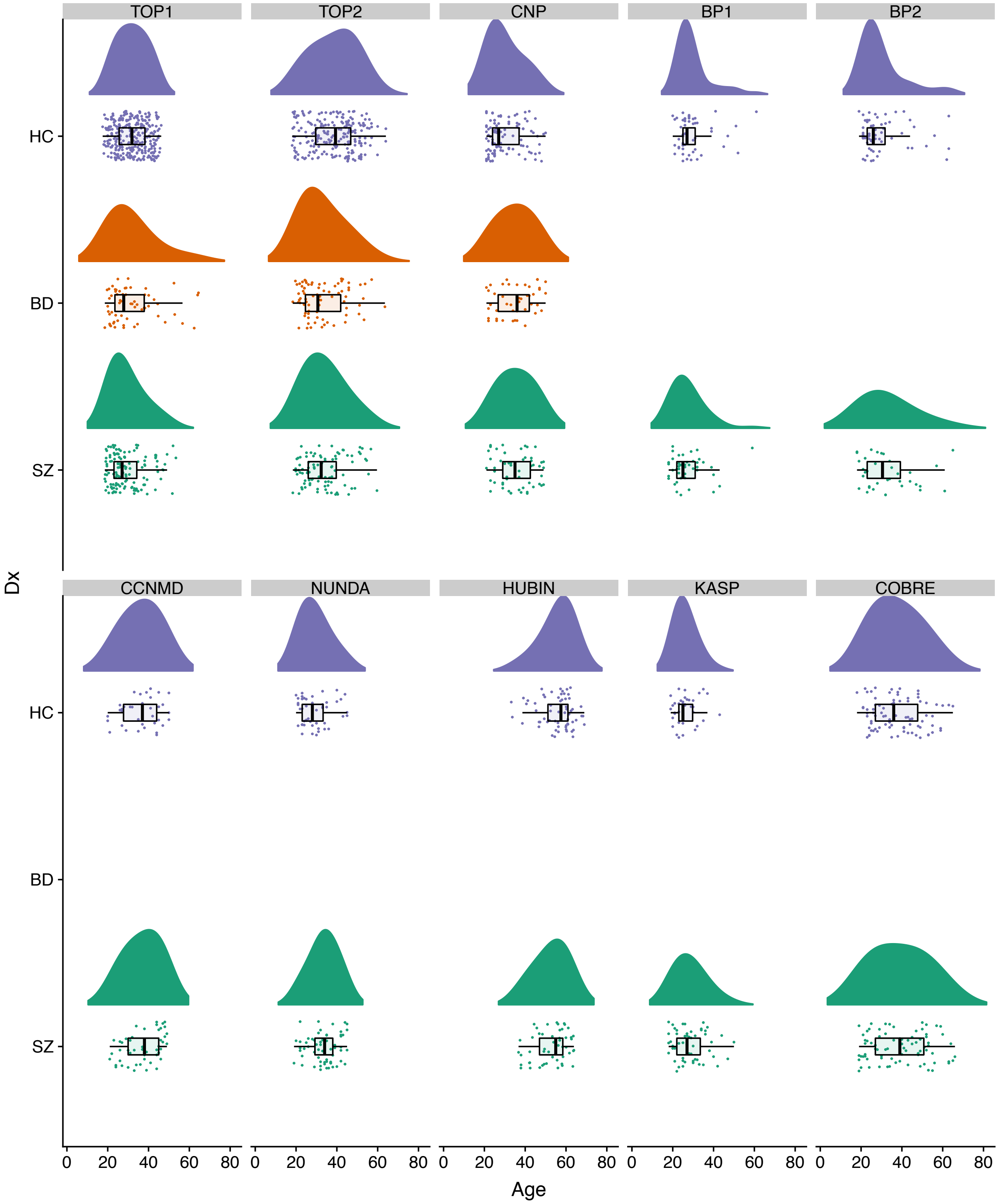


**Supplemental Figure 1.** Raincloud plot depicting the age distribution for each group within each cohort in the test set. Density plots are shown on top with data points and boxplot underneath.

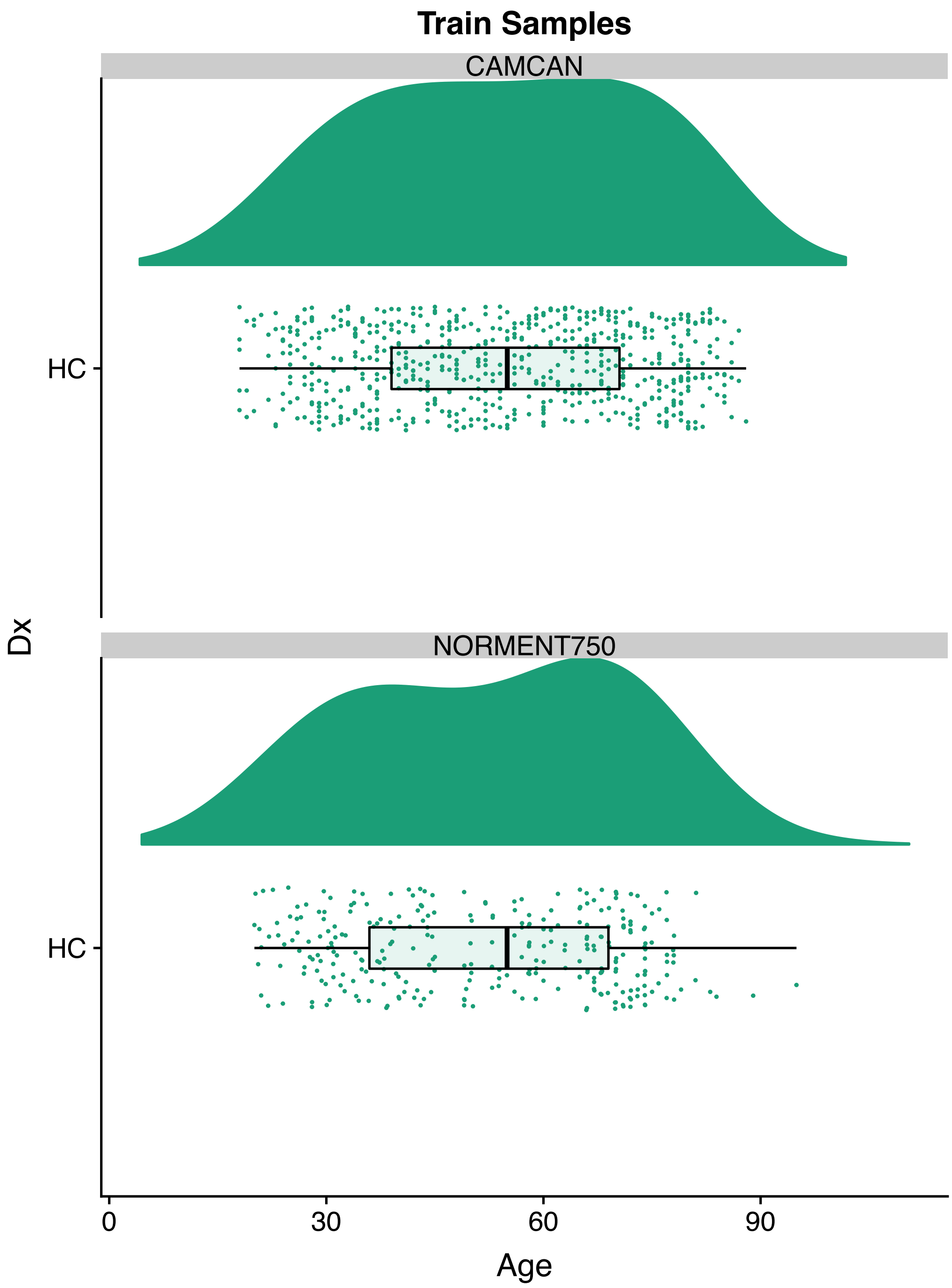


**Supplemental Figure 2.** Raincloud plot depicting the age distribution within the training set. Density plots are shown on top with data points and boxplot underneath.

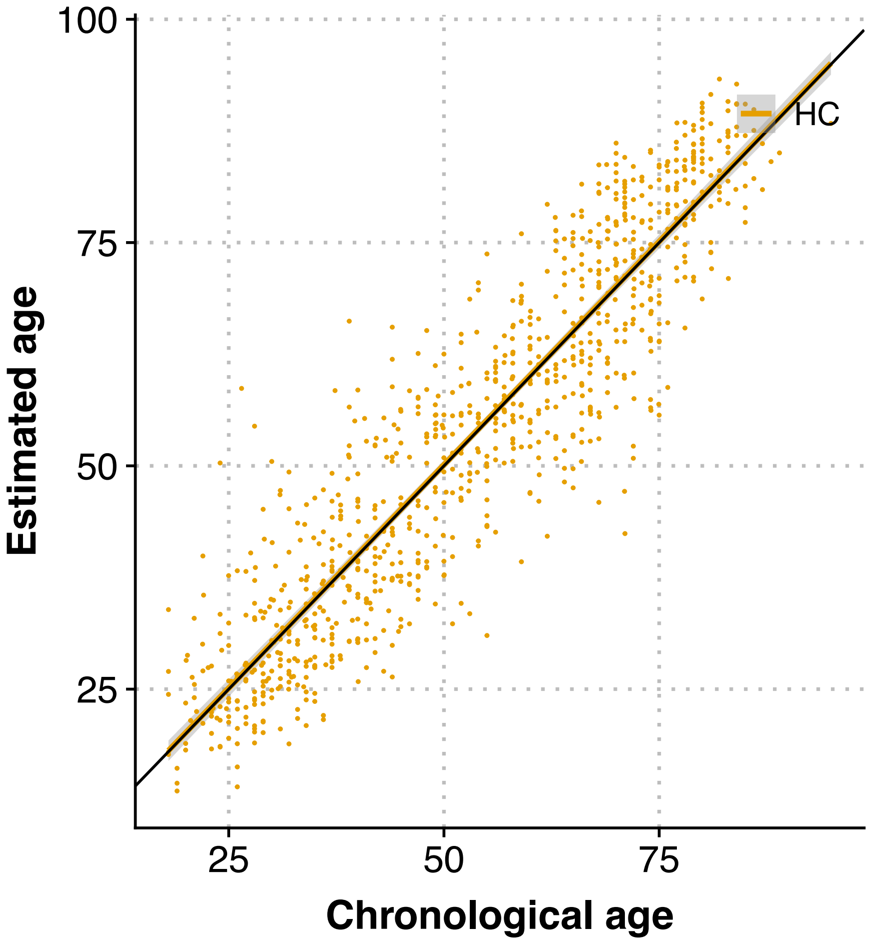


**Supplemental Figure 3.** Predicted age (corrected for age and scanner) based on the all features model plotted as function of chronological age in the training set. The fit lines represent the best linear fit (*r*=.924, 95% CI: .912-.935, MAE=6.49, RMSE=8.08).

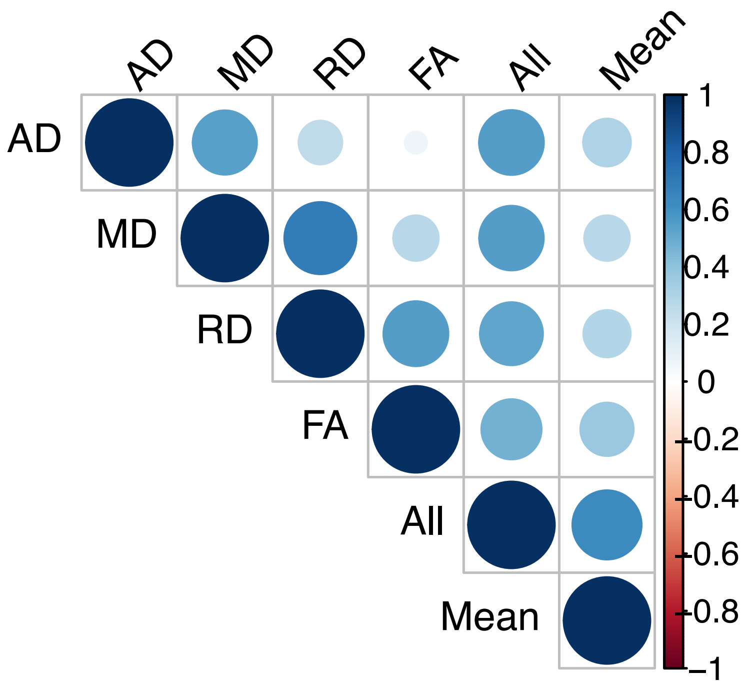


**Supplemental Figure 4.** Correlation matrix showing the relationship between the brain age models.

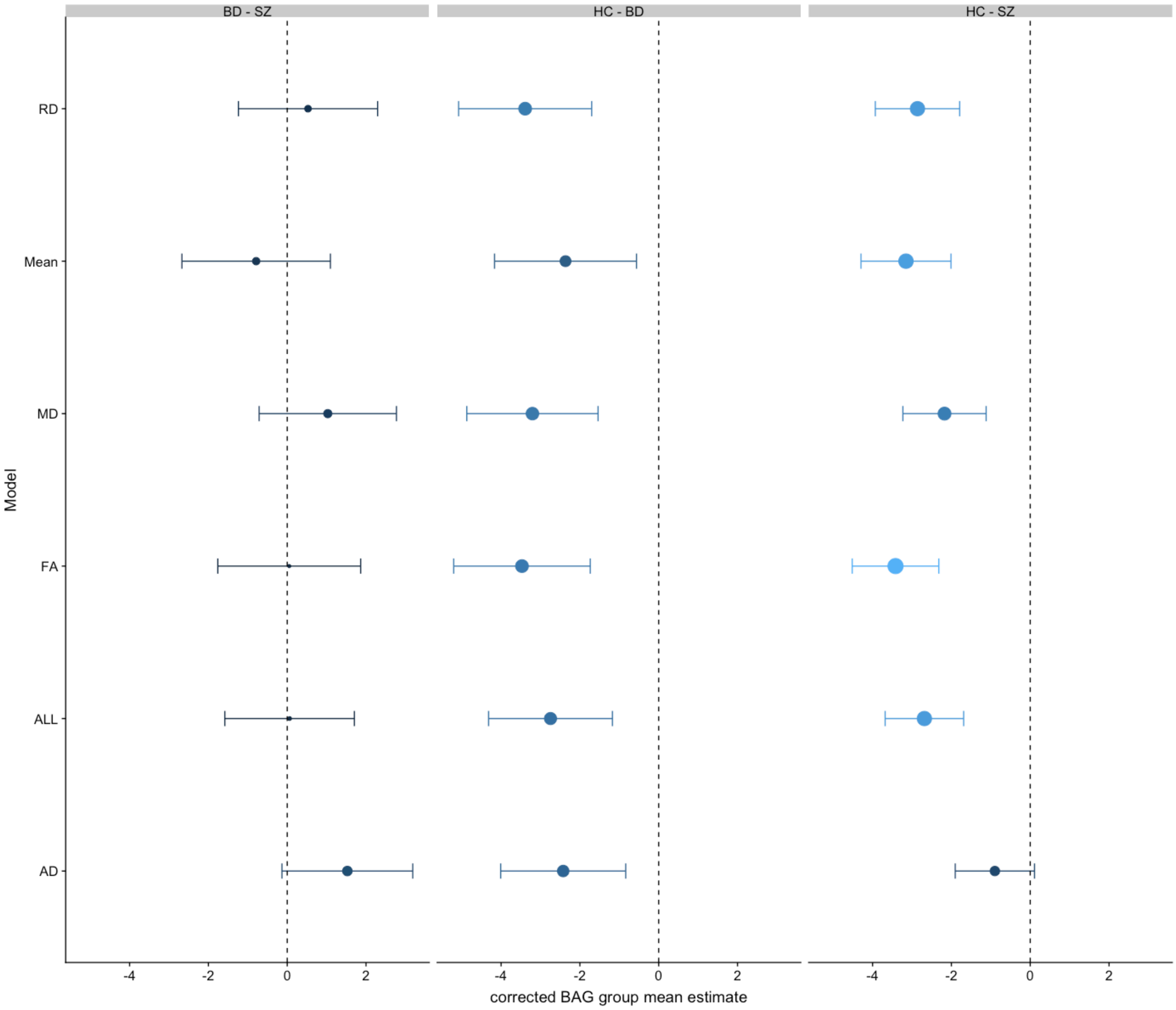


**Supplemental Figure 5.** The figure shows the difference in estimated marginal means in corrected BAG and corresponding 95% confidence interval for the three-group contrast in each model.

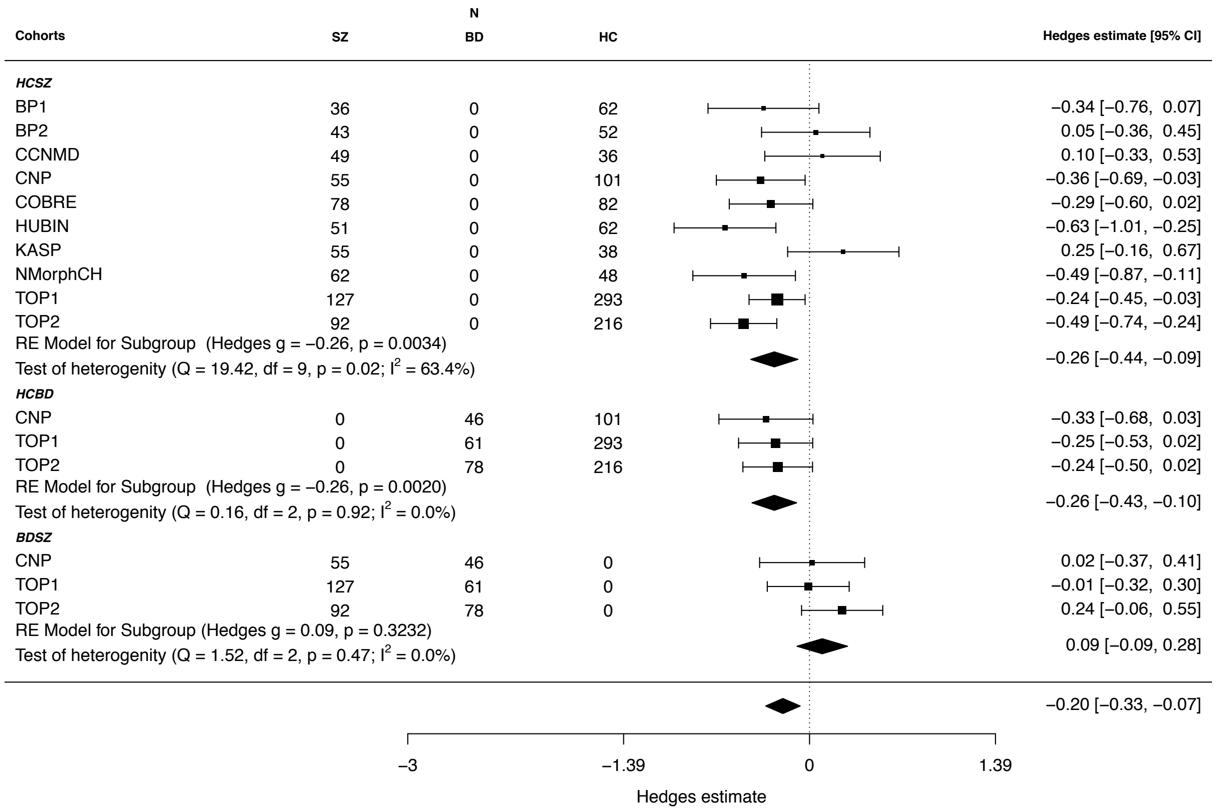


**Supplemental Figure 6.** Forest plot summarizing the results from the meta-analytical approach for the FA features model. Hedges estimate was used to calculate the effect size.

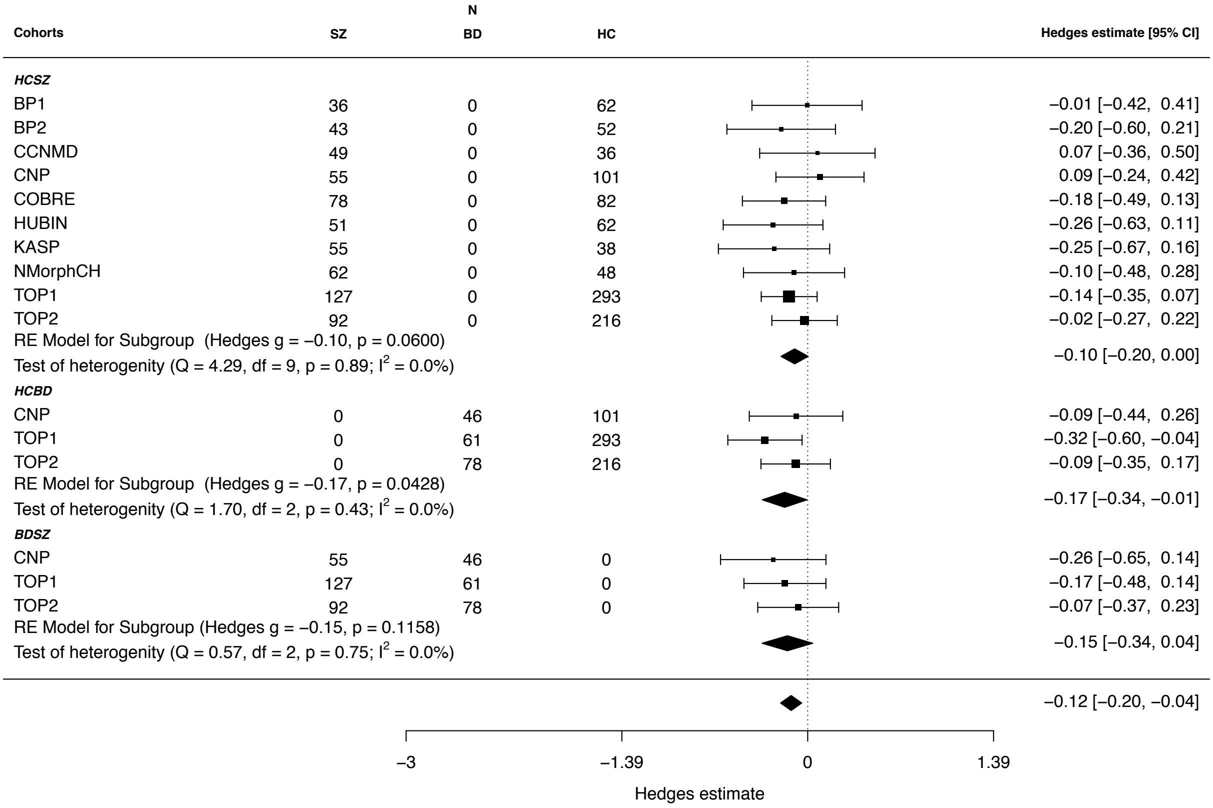


**Supplemental Figure 7.** Forest plot summarizing the results from the meta-analytical approach for the AD features model. Hedges estimate was used to calculate the effect size.

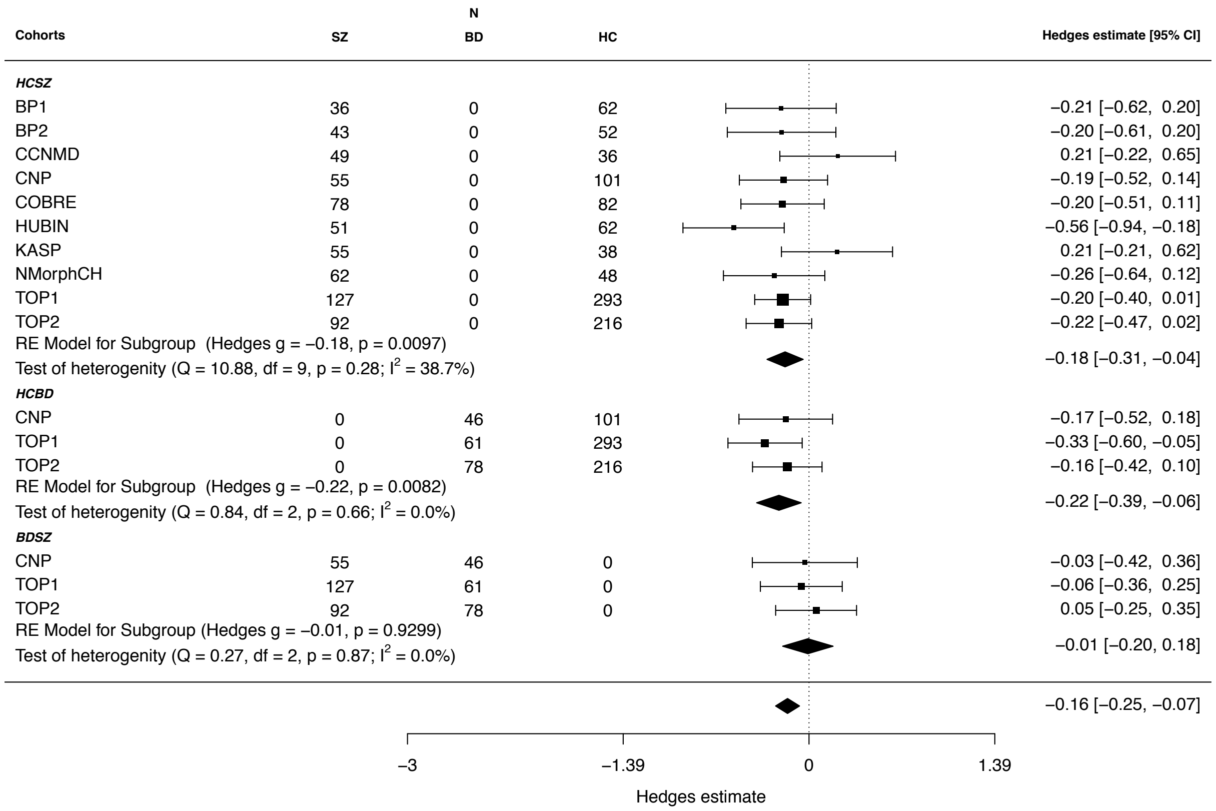


**Supplemental Figure 8.** Forest plot summarizing the results from the meta-analytical approach for the MD features model. Hedges estimate was used to calculate the effect size.

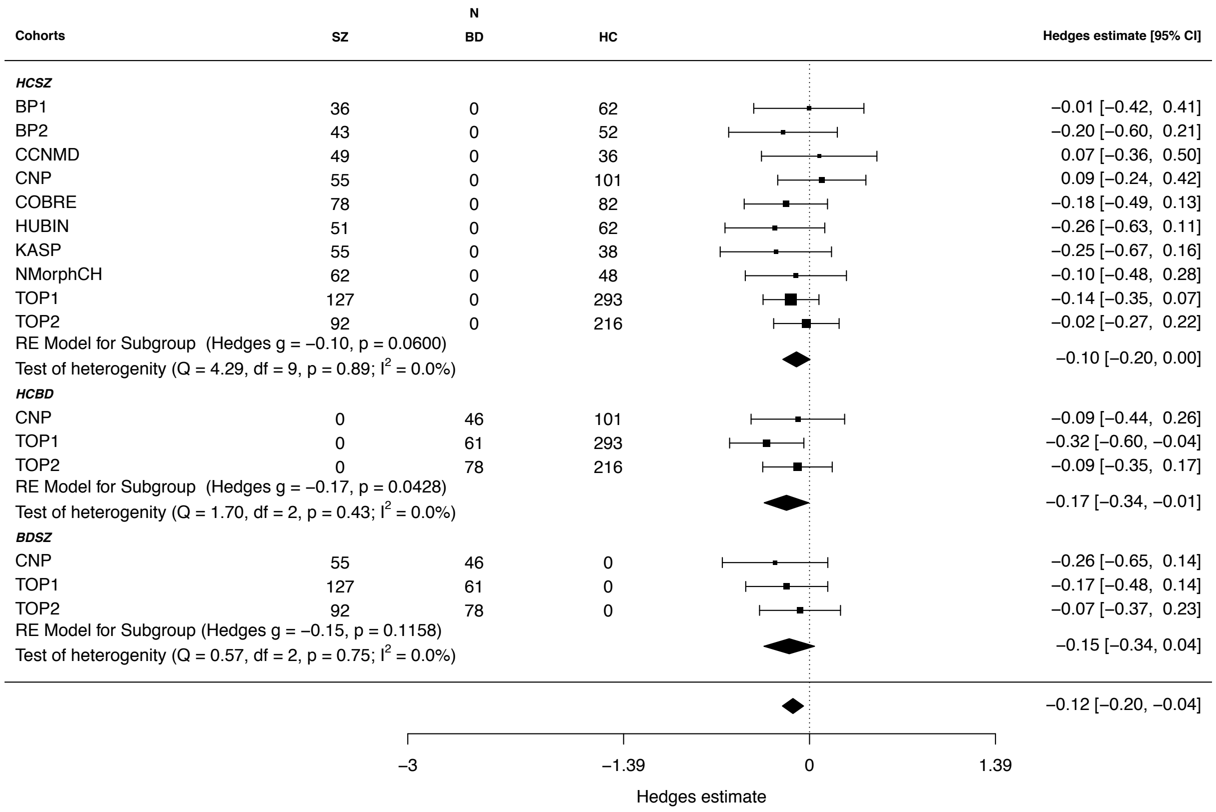


**Supplemental Figure 9.** Forest plot summarizing the results from the meta-analytical approach for the ‘RD features’ model. Hedges estimate was used to calculate the effect size.

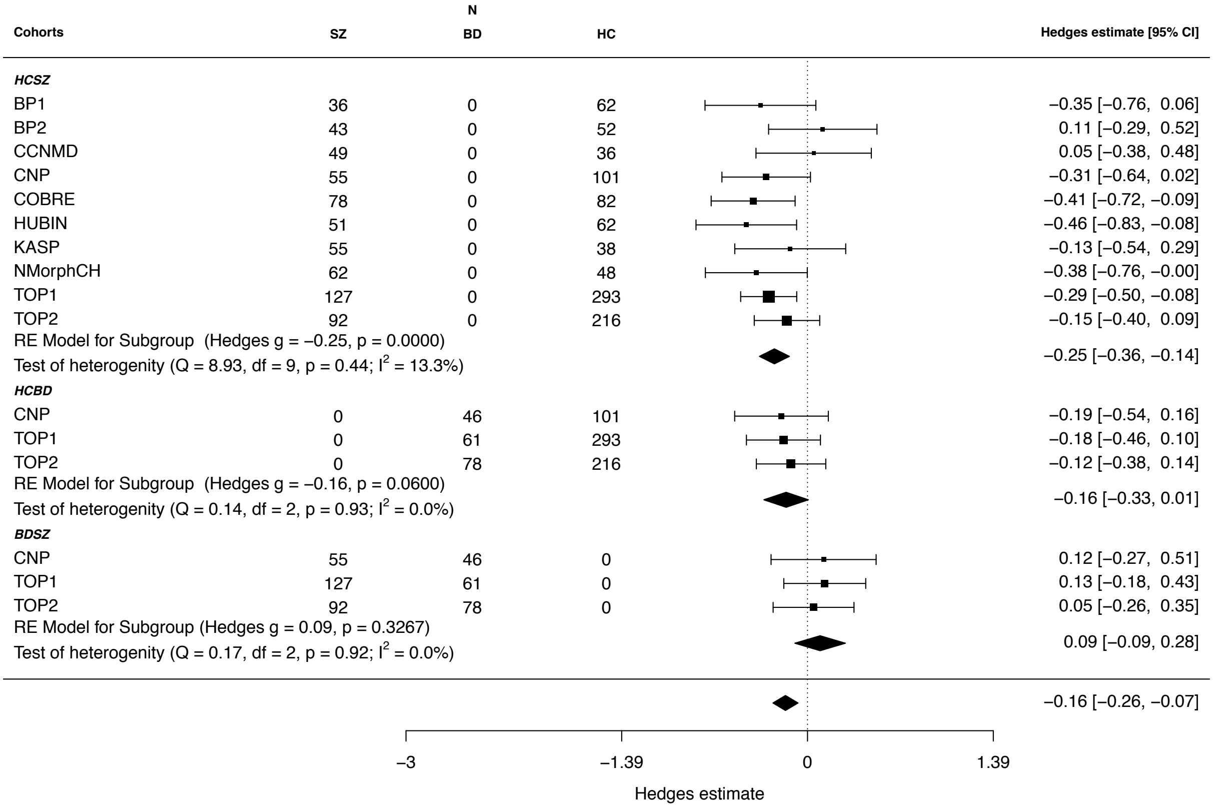


**Supplemental Figure 10.** Forest plot summarizing the results from the meta-analytical approach for the mean skeleton features model. Hedges estimate was used to calculate the effect size.

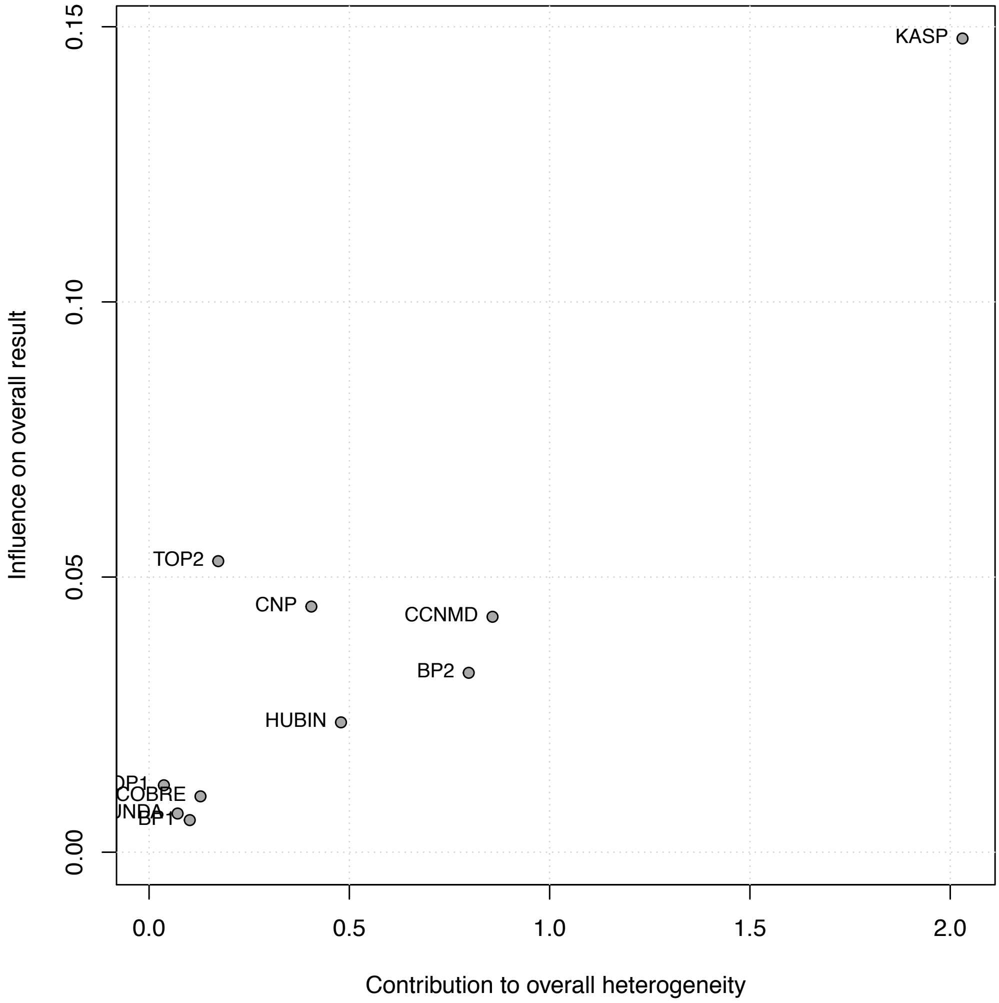


**Supplemental Figure 11.** Baujat plot showing each cohort's influence on the overall effect size and contribution to overall heterogeneity for the HC-SZ contrast. The horizontal axis depicts each study's contribution to the overall heterogeneity, as measured by Cochrans Q. The vertical axis shows the study's influence on the pooled effect size.

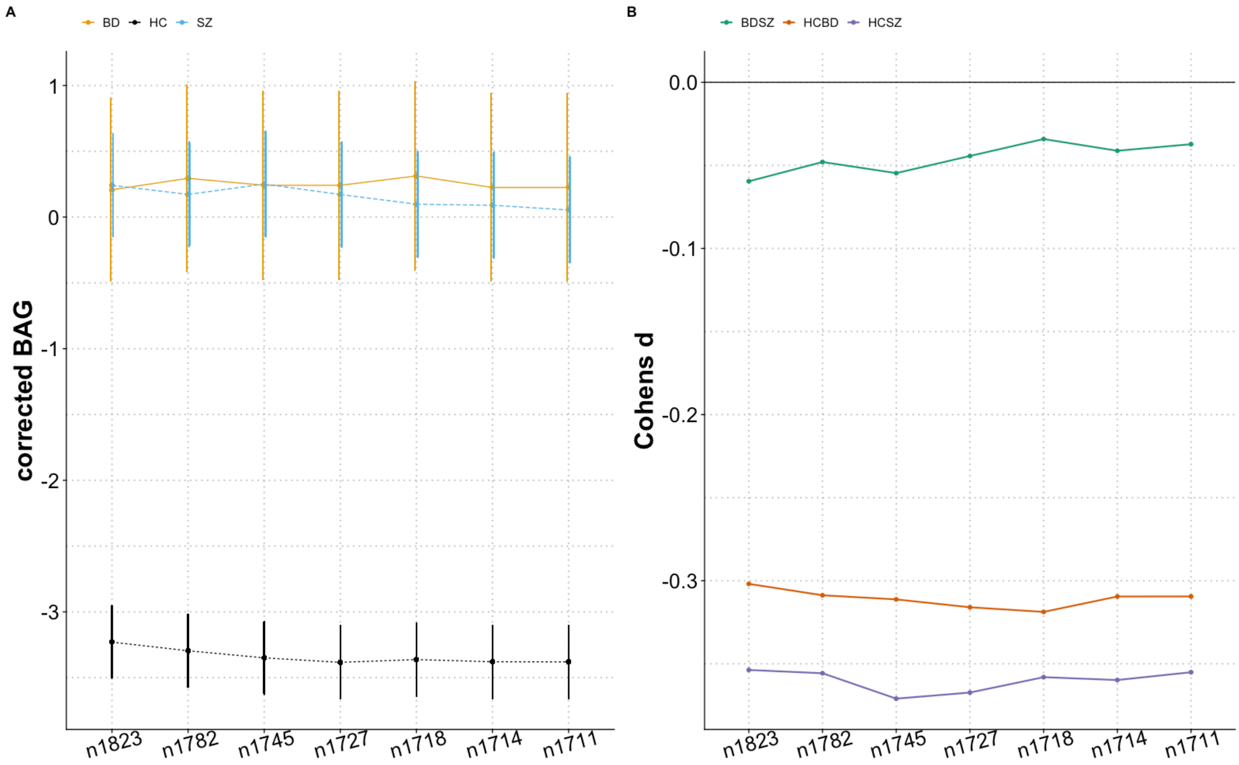


**Supplemental Figure 12.** Quality control using FA-based brain age prediction. A) Residualized brain age gaps plotted as a function of QC stringency/level. B) Relevant effect sizes from group comparisons plotted as a function of QC stringency/level.

**Supplemental Table 1.** Demographics of included cohorts

| Cohort | Cohort N | Number of subjects per group | Age in years: mean ± sd \| range | Sex: m/f |
|---|---|---|---|---|
| Test samples | 1823 | 648 (SZ) | 34.50±11.40 \| 18-66 | 440/208 |
| | | 185 (BD) | 33.12±10.53 \| 18-64 | 89/96 |
| | | 990 (HC) | 34.70 ±11.24 \| 18-68 | 550/440 |
| BP2 | 98 | 36(SZ) | 32.83±12.42 \| 18-65 | 28/8 |
| | | 62(HC) | 30.00±10.88 \| 19-63 | 30/23 |
| BP1 | 95 | 43(SZ) | 27.19±8.07 \| 18-59 | 28/15 |
| | | 52(HC) | 29.69±8.50 \| 20-61 | 35/17 |
| CCNMD | 85 | 49 (SZ) | 37.12 ±8.57 \| 21-49 | 32/17 |
| | | 36 (HC) | 36.28 ±8.94 \| 20-50 | 17/19 |
| COBRE | 161 | 78 (SZ) | 39 ±13.75 \| 18-66 | 62/16 |
| | | 83 (HC) | 37.8 ±11.81 \| 18-65 | 62/21 |
| CNP | 202 | 55 (SZ) | 35.33 ±8.50 \| 21-49 | 42/13 |
| | | 46(BD) | 35.02 ±8.88 \| 21-50 | 26/20 |
| | | 101 (HC) | 30.36 ±8.29 \| 21-50 | 50/51 |
| HUBIN | 113 | 51 (SZ) | 52.55 ±8.01 \| 33-68 | 40/11 |
| | | 62 (HC) | 55.59 ±8.26 \| 33-68 | 41/21 |
| KASP | 93 | 55 (SZ) | 28.47 ±7.69 \| 18-50 | 37/18 |
| | | 38 (HC) | 26.21 ±5.44 \| 19-43 | 22/16 |
| NMorphCH | 110 | 62 (SZ) | 33.56 ±6.93 \| 19-45 | 41/21 |
| | | 48 (HC) | 29.35 ±7.11 \| 20-45 | 23/25 |
| TOP2 | 386 | 92 (SZ) | 33.95 ±10.08 \| 18-59 | 53/39 |
| | | 78 (BD) | 33.08 ±10.52 \| 18-63 | 34/44 |
| | | 216 (HC) | 38.37±11.26 \| 18-64 | 99/117 |
| TOP1 | 481 | 127 (SZ) | 29.48 ±8.55 \| 18-53 | 77/50 |
| | | 61 (BD) | 31.74 ±11.58 \| 18-64 | 29/32 |
| | | 293 (HC) | 31.91 ±7.48 \| 17-46 | 172/121 |
| Train samples | 927 | 927 (HC) | 53.82 ± 18.38 \| 18-94 | 446/481 |
| CAMCAN | 631 | 631 (HC) | 54.43±18.53 \| 18-88 | 313/318 |
| TOP2 | 296 | 296 (HC) | 52.50±17.99 \| 20-94 | 133/163 |

**Supplemental Table 2.** Description of included cohorts

| Cohort | Source | Comment |
|---|---|---|
| CAMCAN | https://camcan-archive.mrc-cbu.cam.ac.uk/dataaccess/ | Data collection and sharing for this project was provided by the Cambridge Centre for Ageing and Neuroscience (CamCAN). CamCAN funding was provided by the UK Biotechnology and Biological Sciences Research Council (grant number BB/H008217/1), together with support from the UK Medical Research Council and University of Cambridge, UK. |
| CCNMD | | CCNMD was supported through NIH Grants P50 MH071616 and R01 MH56584. Reference: Arnedo, Mamah, Baranger, Harms, Barch, Svrakic, et al. (2015): Decomposition of brain diffusion imaging data uncovers latent schizophrenias with distinct patterns of white matter anisotropy. NeuroImage. 120:43-54. |
| DS000030 (CNP) | https://openfmri.org/ | DS000030 work was supported by the Consortium for Neuropsychiatric Phenomics (NIH Roadmap for Medical Research grants UL1-DE019580, RL1MH083268, RL1MH083269, RL1DA024853, RL1MH083270, RL1LM009833, PL1MH083271, and PL1NS062410). Reference: Poldrack, Congdon, Triplett, Gorgolewski, Karlsgodt, Mumford, et al. (2016): A phenome-wide examination of neural and cognitive function. *Scientific data*. 3:160110. |
| COBRE | http://schizconnect.org/ | Data was downloaded from the COllaborative Informatics and Neuroimaging Suite Data Exchange tool (COINS; http://coins.mrn.org/dx) and data collection was performed at the Mind Research Network, and funded by a Center of Biomedical Research Excellence (COBRE) grant 5P20RR021938/P20GM103472 from the NIH to Dr. Vince Calhoun Reference: Cetin, Christensen, Abbott, Stephen, Mayer, Canive, et al. (2014): Thalamus and posterior temporal lobe show greater inter-network connectivity at rest and across sensory paradigms in schizophrenia. NeuroImage. 97:117-126. |
| BERGENPSYKOSE (BP1, BP2) | | European Research Council (ERC AdG) (693124), the Research Council of Norway (213727), and Western Norway Health-Authorities (912045, 911820, 911679) |

| | | |
|---|---|---|
| HUBIN | | This study was supported by the Swedish Research Council (2006- 2992, 2006-986, K2007-62X-15077-04-1, 2008-2167, K2008-62P- 20597-01-3. K2010-62X-15078-07-2, K2012-61X-15078-09-3, 2017-00949, K2015-62X-15077-12-3), the regional agreement on medical training and clinical research between Stockholm County Council and the Karolinska Institutet, the Knut and Alice Wallenberg Foundation, and the HUBIN project. |
| KASP | | KaSP was supported by grants from the Swedish Medical Research Council (SE: 2009-7053; 2013-2838; SC: 523-2014-3467), the Swedish Brain Foundation, Åhlén-siftelsen, Svenska Läkaresällskapet, Petrus och Augusta Hedlunds Stiftelse, Torsten Söderbergs Stiftelse, the AstraZeneca-Karolinska Institutet Joint Research Program in Translational Science, Söderbergs Königska Stiftelse, Professor Bror Gadelius Minne, Knut och Alice Wallenbergs stiftelse, Stockholm County Council (ALF and PPG), Centre for Psychiatry Research, KID-funding from the Karolinska Institutet. |
| NMorphCH | http://schizconnect.org/ | Data used in preparation of this article were obtained from the Neuromorphometry by Computer Algorithm Chicago (NMorphCH) dataset (http://nunda.northwestern.edu/nunda/data/projects/NMorphCH). The investigators within NMorphCH contributed to the design and implementation of NMorphCH and/or provided data but did not participate in analysis or writing of this report. Data collection and sharing for this project was funded by NIMH grantA R01 MH056584 |
| STROKEMRI (TOP2) | | Supported by the Research Council of Norway (249795, 248238), the South-Eastern Norway Regional Health Authority (2014097, 2015044, 2015073, 2016083), and the Norwegian ExtraFoundation for Health and Rehabilitation (2015/FO5146). |
| Sikkerhets-TOP (TOP2) | | South-Eastern Norway Regional Health Authority (2016044) |
| TOP/NORMENT (TOP1, TOP2) | | The work was funded by the Research Council of Norway (213700, 213837, 223273, 204966, 213694, 229129, 249795/F20, 248778), the South-Eastern Norway Regional Health Authority (2013123, 2014097, 2015073, 2017112) and Stiftelsen Kristian Gerhard Jebsen. |

**Supplemental Table 3.** Summary of MRI system and dMRI acquisition for each cohort

| Cohort | Field strength | Scanner | b-value | TR (ms) | TE (ms) | N gradient directions | Voxel resolution |
|---|---|---|---|---|---|---|---|
| BP2 | 3T | GE Discovery MR750 | 1000 | 14000 | 84 | 30 | 1.7 * 1.7 * 2.4 |
| BP1 | 3T | GE Signa HDxt | 1000 | 14000 | 84 | 30 | 1.7 * 1.7 * 2.4 |
| CAMCAN | 3T | Siemens Trio | 1000 | 9100 | 104 | 30 | 2.0 * 2.0 * 2.0 |
| CCNMD | 3T | Siemens Trio | 800 | 8000 | 86 | 30 | 2.0 * 2.0 * 2.0 |
| COBRE | 3T | Siemens Trio | 800 | 9000 | 84 | 30 | 2.0 * 2.0 * 2.0 |
| CNP | 3T | Siemens Trio | 1000 | 9000 | 93 | 64 | 2.0 * 2.0 * 2.0 |
| HUBIN/KASP | 3T | GE Discovery MR750 | 1000 | 6000 | 82 | 60 | 0.9 * 0.9 * 2.0 |
| NMorphCH | 3T | Siemens Trio | 800 | 8000 | 86 | 30 | 2.0 * 2.0 * 2.0 |
| TOP2 | 3T | GE Discovery MR750 | 1000 | 8150 | 83 | 60 | 2.0 * 2.0 * 2.0 |
| TOP1 | 3T | GE Signa HDxt | 1000 | 15000 | 81 | 30 | 1.9 * 1.9 * 2.5 |

**Supplemental Table 4.** Overview of TOI/ROI features

| |
|---|
| Mean skeletonFA |
| Mean skeleton MD |
| Mean skeleton RD |
| Mean skeleton AD |
| Genu of corpus callosum |
| Body of corpus callosum |
| Splenium of corpus callosum |
| Anterior thalamic radiation L |
| Anterior thalamic radiation R |
| Corticospinal tract L |
| Corticospinal tract R |
| Cingulum (cingulate gyrus) L |
| Cingulum (cingulate gyrus) R |
| Cingulum (hippocampus) L |
| Cingulum (hippocampus) R |
| Forceps major |
| Forceps minor |
| Inferior fronto-occipital fasciculus L |
| Inferior fronto-occipital fasciculus R |
| Inferior longitudinal fasciculus L |
| Inferior longitudinal fasciculus R |
| Superior longitudinal fasciculus L |
| Superior longitudinal fasciculus R |
| Uncinate fasciculus L |
| Uncinate fasciculus R |
| Superior longitudinal fasciculus (temporal part) L |
| Superior longitudinal fasciculus (temporal part) R |

**Supplemental Table 5**. Demographics of excluded participants by group

| Group | N (female/male) | Mean age in years, range (sd) |
|---|---|---|
| HC | 59 (21/38) | 32.87, 18.03-51.81 (10.05) |
| SZ | 39 (12/27) | 30.24, 18.60 - 57.76 (9.37) |
| BD | 28 (20/8) | 31.30, 19.31 - 49.00 (8.14) |

The remaining 34 datasets were excluded due to missing key demographic and/or clinical information (such as case/control status).
